# Child abuse associates with increased recruitment of perineuronal nets in the ventromedial prefrontal cortex: a possible implication of oligodendrocyte progenitor cells

**DOI:** 10.1101/2020.10.19.345355

**Authors:** Arnaud Tanti, Claudia Belliveau, Corina Nagy, Malosree Maitra, Fanny Denux, Kelly Perlman, Frank Chen, Refilwe Mpai, Candice Canonne, Stéphanie Théberge, Ashley McFarquhar, Maria Antonietta Davoli, Catherine Belzung, Gustavo Turecki, Naguib Mechawar

**Author notes:** **Corresponding authors: Naguib Mechawar, PhD**, McGill Group for Suicide Studies, Douglas Mental Health Institute, Department of Psychiatry, McGill University, **Arnaud Tanti, PhD**, INSERM UMR 1253, Tours, France. equal contribution (Arnaud Tanti, Claudia Belliveau).

## Abstract

Child abuse (CA) is a strong predictor of psychopathologies and suicide, altering normal trajectories of brain development in areas closely linked to emotional responses such as the prefrontal cortex (PFC). Yet, the cellular underpinnings of these enduring effects are unclear. Childhood and adolescence are marked by the protracted formation of perineuronal nets (PNNs), which orchestrate the closure of developmental windows of cortical plasticity by regulating the functional integration of parvalbumin interneurons (PV) into neuronal circuits. Using well-characterized post-mortem brain samples, we show that a history of CA is specifically associated with increased densities and morphological complexity of WFL-labeled PNNs in the ventromedial PFC (BA11/12), possibly suggesting increased recruitment and maturation of PNNs. Through single-nucleus sequencing and fluorescent in-situ hybridization, we found that the expression of canonical components of PNNs is enriched in oligodendrocyte progenitor cells (OPCs), and that they are upregulated in CA victims. These correlational findings suggest that early-life adversity may lead to persistent patterns of maladaptive behaviors by reducing the neuroplasticity of cortical circuits through the enhancement of developmental OPC-mediated PNN formation.

## Introduction

Child abuse (CA) has enduring effects on psychological development. Severe adversity during sensitive periods, during which personality traits, attachment patterns, cognitive functions and emotional responses are shaped by environmental experiences, has a profound effect on the structural and functional organization of the brain ^1^.

At the cellular level, childhood and adolescence are marked by the protracted maturation of neural circuits, characterized by windows of heightened plasticity that precede the development of functional inhibitory connections and the balance of excitatory-inhibitory neurotransmission ^2^. A major mechanism involved in this process is the recruitment of perineuronal nets (PNNs), a condensed form of extracellular matrix forming most notably around parvalbumin-expressing (PV+) interneurons. PNNs are thought to gradually decrease heightened plasticity by stabilizing the integration and function of PV+ cells into cortical networks and hindering the remodeling of these networks ^3,4^. This has been notably linked to the persistence of long-term associations, including fear memories ^5–7^.

Evidence in rodents suggests that early-life stress associates with precocious functional maturation of PV+ neurons and the early emergence of adult-like characteristics of fear and extinction learning ^8^, in addition to discrete changes in the immunoreactivity of inhibitory neuron markers and PNNs ^9^. Taken together, these observations suggest that CA may alter the formation of PNNs.

We addressed this question using well-characterized post-mortem samples (Douglas-Bell Canada Brain Bank) from adult depressed suicides who died during an episode of major depression with (DS-CA) or without (DS) a history of severe CA and from matched psychiatrically healthy individuals (CTRL). Standardized psychological autopsies were conducted to provide comprehensive post-mortem diagnosis and retrieve various dimensions of childhood experience, including history and severity of CA. We focused on the ventromedial prefrontal cortex (vmPFC), encompassing Brodmann areas 11 and 12 in our study, a brain area closely linked to emotional learning and which is structurally and functionally altered in individuals with a history of CA ^1,10–13^.

## Materials and Methods

### Human post-mortem brain samples

Brain samples were obtained from the Douglas-Bell Canada Brain Bank (Montreal, Canada). Phenotypic information was retrieved through standardized psychological autopsies, in collaboration with the Quebec Coroner’s Office and with informed consent from next of kin. Presence of any or suspected neurological/neurodegenerative disorder signaled in clinical files constituted an exclusion criterion. Cases and controls are defined with the support of medical charts and Coroner records. Proxy-based interviews with one or more informants best acquainted with the deceased are supplemented with information from archival material obtained from hospitals, Coroner’s office and social services. Clinical vignettes are then produced and assessed by a panel of clinicians to generate DSM-IV diagnostic criteria, providing sociodemographic characteristics, social developmental history, DSM-IV axis I diagnostic information and behavioural traits; information that is obtained through different adapted questionnaires. Toxicological assessments and medication prescription are also obtained. As described previously^14^, characterization of early-life histories was based on adapted Childhood Experience of Care and Abuse (CECA) interviews assessing experiences of sexual and physical abuse, as well as neglect^15^, and for which scores from siblings are highly concordant^16^. We considered as severe early-life adversity reports of non-random major physical and/or sexual abuse during childhood (up to 15 years). Only cases with the maximum severity ratings of 1 and 2 were included. This information was then complemented with medical charts and coroner records. Because of this narrow selection criterion, it was not possible to stratify different types of abuse within the sample.

Subject characteristics are described in **Table 1**. Correlations between covariates (age, post-mortem interval (PMI), pH, substance dependence and medication) and the variables measured in our study are presented in **Supplementary Table 1**.

**Table 1.**
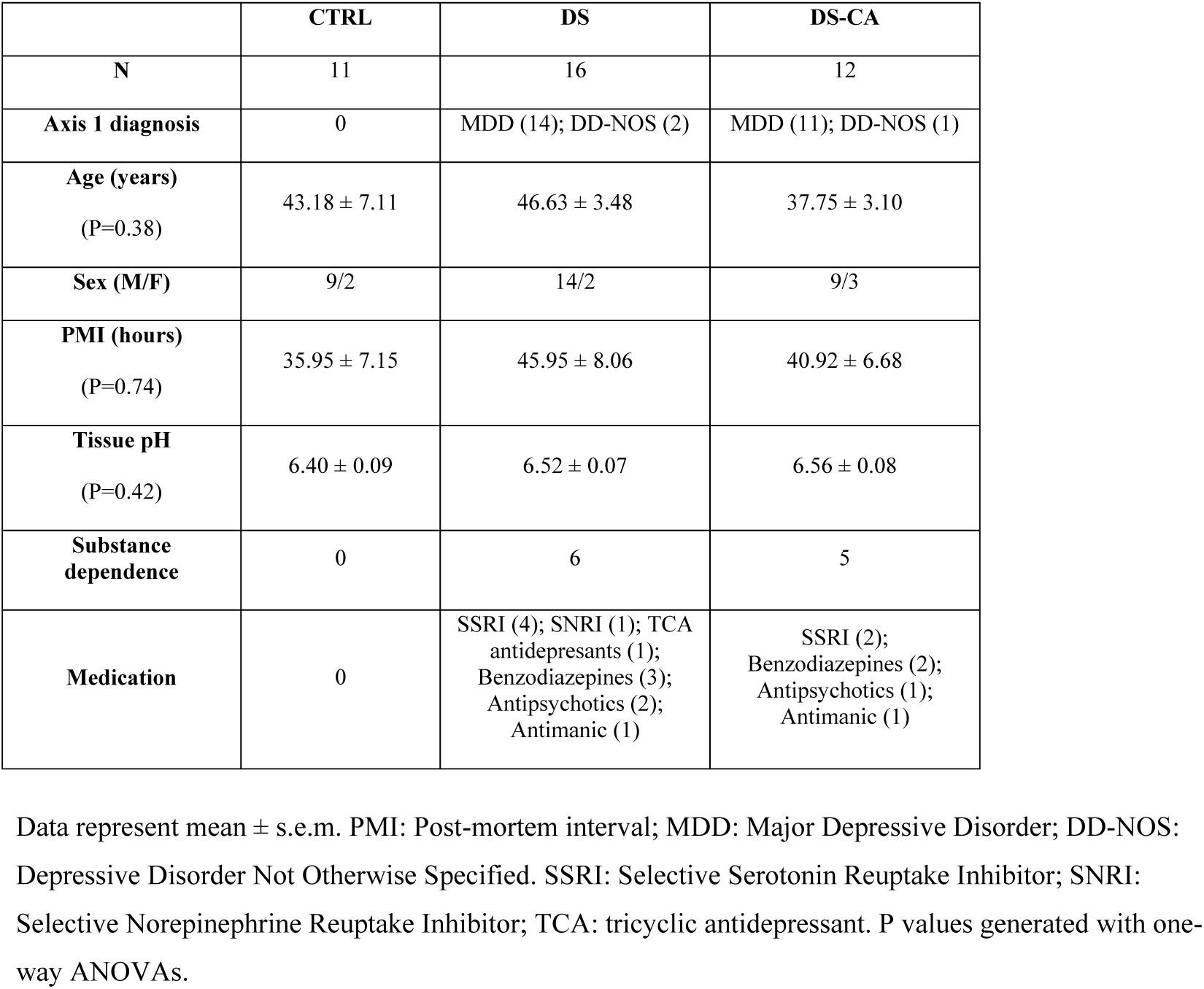
Group characteristics.

|  | **CTRL** | **DS** | **DS-CA** |
| --- | --- | --- | --- |
| **N** | 11 | 16 | 12 |
| **Axis 1 diagnosis** | 0 | MDD (14); DD-NOS (2) | MDD (11); DD-NOS (1) |
| **Age (years)**<br>(P=0.38) | 43.18 ± 7.11 | 46.63 ± 3.48 | 37.75 ± 3.10 |
| **Sex (M/F)** | 9/2 | 14/2 | 9/3 |
| **PMI (hours)**<br>(P=0.74) | 35.95 ± 7.15 | 45.95 ± 8.06 | 40.92 ± 6.68 |
| **Tissue pH**<br>(P=0.42) | 6.40 ± 0.09 | 6.52 ± 0.07 | 6.56 ± 0.08 |
| **Substance dependence** | 0 | 6 | 5 |
| **Medication** | 0 | SSRI (4); SNRI (1); TCA antidepressants (1); Benzodiazepines (3); Antipsychotics (2); Antimanic (1) | SSRI (2); Benzodiazepines (2); Antipsychotics (1); Antimanic (1) |

Data represent mean ± s.e.m. PMI: Post-mortem interval; MDD: Major Depressive Disorder; DD-NOS: Depressive Disorder Not Otherwise Specified. SSRI: Selective Serotonin Reuptake Inhibitor; SNRI: Selective Norepinephrine Reuptake Inhibitor; TCA: tricyclic antidepressant. P values generated with one-way ANOVAs.

### Tissue dissections

Dissections were performed by expert brain bank staff on fresh-frozen 0.5 cm thick coronal sections with the guidance of a human brain atlas^17^. Ventromedial prefrontal cortex samples were dissected in sections equivalent to plate 3 (approximately − 48 mm from the center of the anterior commissure) of this atlas, corresponding to Brodmann areas 11 and 12. Samples were either kept frozen or fixed overnight in 10% formalin until processed for in situ hybridization or immunofluorescence, respectively. Samples used for PV immunohistochemistry were stored long term in 10% formalin until processed.

### Immunohistochemistry

Frozen tissue blocks were fixed in 10% neutral buffered formalin overnight at 4°C, rinsed in PBS and kept in 20% sucrose/PBS solution until serially sectioned at 40 μm on a cryostat. Free-floating sections were rinsed in phosphate-buffered saline (PBS) and then incubated overnight at 4ºC under constant agitation with the antibody (mouse anti-NeuN (Millipore, 1:500, MAB377), goat anti-Versican (R&D, 1:100, AF3054)) or lectin (biotinylated Wisteria Floribunda Lectin [WFL], Vector Laboratories, B-1355, 1:2500) of interest diluted in a blocking solution of PBS/0.2% Triton-X/2% normal donkey serum. Sections were then rinsed and incubated for 2h at room temperature with the appropriate fluorophore conjugated secondary antibody (Alexa-488 anti-Mouse (Jackson ImmunoResearch, 1:500) for NeuN, Dylight-594 anti-goat (Jackson ImmunoResearch, 1:500) for VCAN, or Cy3-conjugated Streptavidin (Jackson ImmunoResearch, 016-160-084; 1:500) for the detection of PNNs and, diluted in the same blocking solution as the primary incubation. Next, sections were rinsed and endogenous autofluorescence from lipofuscin and cellular debris was quenched with Trueblack (Biotium), omitted for tissues used for intensity measurements. Sections were mounted on Superfrost charged slides and coverslipped with Vectashield mounting medium (Vector Laboratories, H-1800).

Whole vmPFC sections were scanned on a Zeiss Axio Imager M2 microscope equipped with a motorized stage and Axiocam MRm camera at 20x. The ImageJ^18^ software (NIH) Cell Counter plugin was used by a blinded researcher to manually count PNNs. An average of 4 sections per subject was used. Cortical layers were delineated based on NeuN+ cells distribution and morphology, and the number of PNNs as well as the area of each layer were measured, allowing to generate PNNs density values (n/mm^2^). Densities were obtained by averaging by subject the density of PNNs per layer and section, and then averaging subjects’ densities to yield group means.

For PV immunohistochemistry, tissue blocks stored in 10% formalin were first transferred to 30% sucrose/PBS. Once sunk, blocks were flash frozen in isopentane and stored at -80°C until embedded and serially sectioned on a sliding microtome (40 μm). Prior to immunohistochemical staining, tissues underwent antigen retrieval by incubating for 15 minutes in hot 10mM sodium citrate buffer pH=6.0 (Sigma cat. no. S-4641). Sections were rinsed in PBS and incubated in 3% H_2_O_2_/PBS for 15 minutes. After being rinsed, sections were incubated overnight at 4°C under constant agitation with a mouse anti-Parvalbumin antibody (Swant, 1:500, PV235) diluted in a blocking solution of PBS/0.2% Triton-X/2% normal horse serum. Sections were then rinsed and incubated for 2h at room temperature in biotinylated horse anti-mouse antibody (1:500, Vector Laboratories Inc., BA-2001, Burlington, ON, Canada). Then, sections were incubated in the avidin-biotin complex procedure (ABC Kit, Vectastain Elite, Vector Laboratories Inc., Burlington, ON, Canada) for 30 minutes at room temperature. Labeling was revealed with the diaminobenzidine kit (Vector Laboratories Inc., Burlington, ON, Canada) then sections were rinsed and mounted on Superfrost charged glass slides, dehydrated and coverslipped with Permount (Fisher Scientific Inc., Pittsburgh, PA, USA). Immunohistological controls were performed by omitting primary antibodies. After a first round of imaging, the coverslips were removed, and samples were counterstained with cresyl violet to differentiate cortical layers and imaged a second time. Image acquisition was performed on an Olympus VS120 Slide Scanner at 10x. Image analysis was performed in QuPath^19^ (v 0.1.2). Automatic cell detection was used to detect PV+ cells. DAB images were overlaid on the cresyl counterstained image using the function Interactive Image Alignment – which allowed a blinded researcher to delineate the cortical layers based on cresyl violet stained cells. Densities were calculated by cortical layer.

### Fluorescent in situ hybridization

Frozen vmPFC (BA11/12) blocks were cut serially with a cryostat and 10μm-thick sections collected on Superfrost charged slides. In situ hybridization was performed using Advanced Cell Diagnostics RNAscope® probes and reagents following the manufacturer instructions. Sections were first fixed in cold 10% neutral buffered formalin for 15 minutes, dehydrated by increasing gradient of ethanol bathes and air dried for 5 minutes. Endogenous peroxidase activity was quenched with hydrogen peroxide for 10 minutes followed by protease digestion for 30 min at room temperature (omitted for samples undergoing subsequent WFL staining). The following probes were then hybridized for 2 hours at 40°C in a humidity-controlled oven: Hs-PVALB (cat. no. 422181), Hs-VCAN (cat. no. 430071-C2), Hs-PDGFRA (cat. no. 604481-C3), Hs-TNR (cat. no. 525811), Hs-PTPRZ1 (cat. no. 584781-C2), Hs-SLC17A7 (cat. no. 415611), Hs-GAD1 (cat. no. 573061-C3). Amplifiers were added using the proprietary AMP reagents, and the signal visualized through probe-specific HRP-based detection by tyramide signal amplification with Opal dyes (Opal 520, Opal 570 or Opal 690; Perkin Elmer) diluted 1:700. Slides were then coverslipped with Vectashield mounting medium with DAPI for nuclear staining (Vector Laboratories) and kept at 4°C until imaging. Both positive and negative controls provided by the supplier (ACDbio) were used on separate sections to confirm signal specificity. For immunohistochemical staining of PNNs following PVALB, GAD1, or SLC17A7 in situ hybridization, slides were rinsed in PBS, incubated for 30 minutes at 4°C with biotinylated WFL followed by 488-conjugated Streptavidin for 2 hours prior to coverslipping. To better define the cellular identity of neuronal populations surrounded by WLF-labeled PNNs, TrueBlack (Biotium) was used to remove endogenous autofluorescence from lipofuscin and cellular debris.

### Cellular identity and ratios of each cell type surrounded by WFL-labeled PNNs

Image analysis was performed in QuPath (v 0.2.3). Each subject had two sections stained with various cellular markers: DAPI, PVALB, SLC17A7, GAD1 and WFL. To identify the population of cells covered by PNNs and calculate the percentage of each cell type that is covered by a net; a blinded researcher manually identified PNNs and categorized each nucleus within an ROI (spanning layer 3-6) dependent on the presence of canonical cellular markers. A total of 3145 PNNs were classified and a total of 18600 SLC17A7+, 2209 GAD1+/PVALB+ and 8659 GAD1+/PVALB-cells were classified.

A replication experiment was conducted for the proportions of PVALB+ cells enwrapped by WFL-labeled PNNs, which were determined in a single section with an average of 55 PVALB+ cells per subject imaged under a 20x objective through layers 4-5 of the vmPFC.

### Imaging and analysis of in situ RNA expression in OPCs

Image acquisitions was performed on a FV1200 laser scanning confocal microscope (FV1200) equipped with a motorized stage. For each experiment and subject, 6 to 10 stack images were taken to capture at least 20 OPCs (PDGFRA+) per subject. Images were taken using a 60x objective (NA = 1.42) with a XY pixel width of ∼0.25μm and Z spacing of 0.5μm. Laser power and detection parameters were kept consistent between subjects for each set of experiment. Since TSA amplification with Opal dyes yields a high signal to noise ratio, parameters were optimized so that autofluorescence from lipofuscin and cellular debris was filtered out. OPCs were defined by bright clustered puncta-like PDGFRA signal present within the nucleus of the cells. Using ImageJ, stacks were first converted to Z-projections, and for each image the nuclei of OPCs were manually contoured based on DAPI expression. Expression of VCAN, TNR or PTPRZ1 in OPCs was quantified using the area fraction, whereby for each probe the signal was first manually thresholded by a blinded researcher and then the fraction of the contoured nucleus area covered by signal was measured for each OPC. Area fraction was the preferred measure to reflect RNA expression as punctate labeling generated by FISH often aggregates into clusters that cannot readily be dissociated into single dots or molecules.

### Intensity, area and distance measurements

For each subject, ∼15 z-stacks (0.26μm Z-spacing) spanning all layers of the vmPFC were acquired at 40x magnification on an Olympus FV1200 laser scanning confocal microscope. Images for intensity measurement were all acquired at the same laser strength and voltage to avoid imaging differences in intensity or over-exposure. PNNs were traced manually with ImageJ by a blinded researcher using maximum intensity projections generated from each stack. All the PNNs that were observed were traced as long as their whole morphology was in the field of view. For each PNN, the mean pixel value of adjacent background was subtracted to the mean pixel value of the contoured soma of the PNN, yielding the mean corrected fluorescence intensity. To infer on their morphological complexity, we measured the area covered by each contoured PNN, including soma and ramifications extending over proximal dendrites.

To quantify closest distance between OPCs and PV+ cells, low magnification (10x) images of PDGFRA and PVALB FISH sections were taken by a blinded researcher along layers IV and V of the vmPFC, using the granular layer IV as a visual reference. For each PDGFRA+ cell in the field of view, the distance to the nearest PVALB+ cell was measured using the measure tool in imageJ. An average of 90 OPCs per group were quantified.

### OPCs density measurements

Image acquisition was performed on an Olympus VS120 Slide Scanner at 20x. Image analysis was performed in QuPath (v 0.2.3) by a blinded researcher. Automatic cell detection was used to detect DAPI nuclei. Then an object classifier was trained on 5 training images, from 5 different subjects. Cells were deemed PDGFRA+ based on the mean intensity of PDGFRA staining compared to the mean staining of a background channel. In total, 21 subjects were included (CTRL = 6, DS = 7, DS-CA = 8) in this analysis.

### Cell-type specific expression of PNN canonical components using single-nucleus sequencing

Cell-type specific expression of canonical components of PNNs was explored using a snRNA-seq dataset from the human dorsolateral prefrontal cortex (BA9) previously generated by our group ^20^, for which methodology is extensively described in this published resource. Average expression for each PNN component in each cell type was calculated by weighting the expression values (normalized transcript counts) of each cluster by the size (number of nuclei) of the cluster. Weighted average expression values are displayed in a heatmap, scaled by row (i.e. gene). The color bar therefore represents the expression values as z-scores, with darker colors indicating higher expression.

### Statistical analyses

Independently of PMI, pH and approach used, quality of post-mortem samples is notoriously variable and influenced by tissue degradation, quality of fixation and other artefacts. Only samples that showed reliable labeling were included in the different experiments without prior knowledge about group affiliation. Analyses were performed on Statistica version 12 (StatSoft) and Prism version 6 (GraphPad Software). Distribution and homogeneity of variances were assessed with Shapiro–Wilk and Levene’s tests, respectively. PNNs densities were analyzed using a mixed-effects model, using layer and group as fixed factors, followed by Tukey’s HSD test for corrected post hoc comparisons. For all other variables (WFL intensity, WFL area per PNN, PNN+/PVALB ratios, RNA expression in OPCs and distance of OPCs from PVALB+ cells) group effects were detected using one-way ANOVAs or Kruskal-Wallis test followed by Tukey’s HSD or Dunn’s test respectively. Linear regressions and Spearman’s correlation were used to address the relationship between dependent variables and covariates (age, PMI and pH, medication and substance dependence) (Supplementary Table 1). Statistical tests were two sided. Significance threshold was set at 0.05, and all data presented represent mean ±sem.

## Results

PNN densities, visualized by Wisteria Floribunda Lectin (WFL) labeling and NeuN immunostaining (**Fig.1A**) were markedly higher through layers III to VI of vmPFC (BA11/12) samples from individuals with a history of CA compared to controls and depressed suicides with no history of CA (**Fig.1B)**. Although the recruitment of PNNs is developmentally regulated, we did not find any correlation between age and densities of PNNs (Supplementary Table 1), perhaps suggesting that PNN recruitment may already have reached a plateau in our cohort^21^. Likewise, controlling for age did not affect our results (ANCOVA with group as fixed factor and age as covariate; group effect: F(2,35)=11.230, P<0.001). To investigate if CA also associates with maturational or morphological changes of PNNs, we compared the intensity of WFL staining between groups (**Fig.1C**) as an indication of their maturity, as well as the area covered by individual PNNs as an indicator of their morphological complexity (Fig.1D). CA was both associated with higher intensity of WFL staining per PNN (**Fig.1C**) and cells were more extensively covered by PNNs (**Fig.1D**), suggesting overall that CA may precipitate the maturation and the recruitment of PNNs.

**Figure 1.**
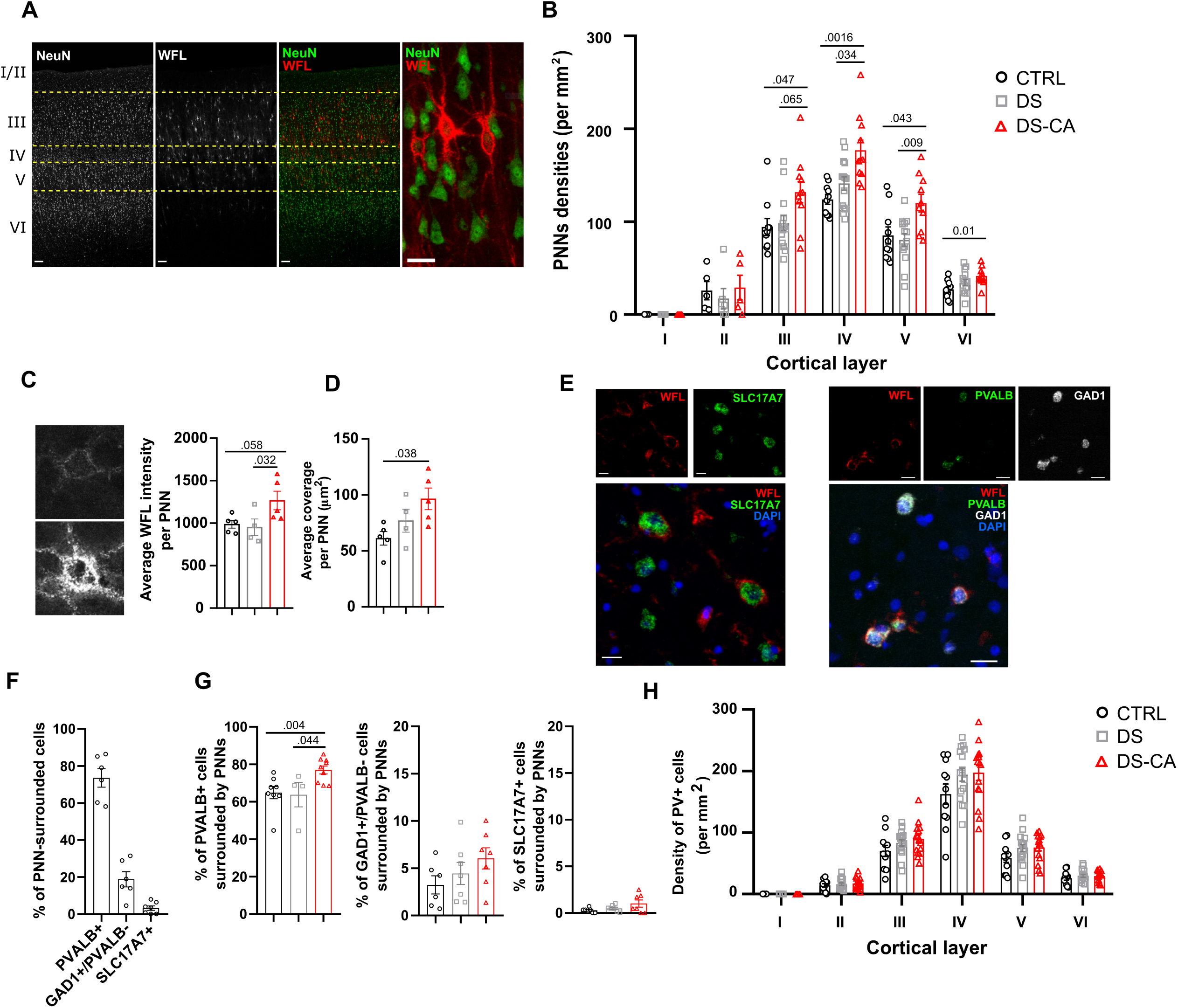
(**A**) Representative images of PNNs labeled with WFL and their distribution throughout human vmPFC cortical layers. Scale bars = 100 and 20 μm (high magnification panel) (**B**) Depressed suicides with a history of child abuse (DS-CA, N = 11) have significantly higher PNN densities compared to controls (CTRL, N = 10) and depressed suicides without history of child abuse (DS, N= 14) (group effect: F (2, 32) = 7.029, P=0.0029; layer effect: F (3.395, 78.09) = 194.2, P<0.0001; layer x group: F (10, 115) = 2.07, P= 0.0029, followed by Tukey’s multiple comparison test). (**C**) Representative images of a low (top) and high (bottom) intensity PNN in the vmPFC. PNNs from DS-CA subjects (N=5) showed higher average WFL intensity compared to CTRLs (N=5) or DS (N=4) (Kruskal-Wallis ANOVA: H(2, 14) = 5.57, P=0.049, followed by Dunn’s test) (**D**) PNNs from DS-CA subjects (N=5) showed higher complexity (area covered by PNNs) compared to CTRLs (N=5) or DS (N=4) (Kruskal-Wallis ANOVA: H (2, 14) = 6.223, P=0.034, followed by Dunn’s test). (**E**) Left: Representative images of in situ hybridization for SLC17A7 (green) followed by WFL labeling (red). Nuclei were stained with DAPI (blue); Right: Representative images of in situ hybridization for PVALB (green) and GAD1 (white) followed by WFL labeling (red). Nuclei were stained with DAPI (blue). Scale bars = 20μm. (**F**) Proportions of WFL-labeled PNNs expressing PVALB (PV neurons), GAD1 but not PVALB (other inhibitory neurons), and SLC17A7 (excitatory neurons). (**G**) DS-CA (N=9) subjects have higher ratios of PVALB+ cells surrounded by PNNs compared to CTRLs (N =8) and DS subjects (N=4) (Kruskal-Wallis ANOVA H(2,21) = 9.45, P=0.0037, followed by Dunn’s test), but not of GAD1+/PVALB-cells (Krukal-Wallis ANOVA H(2,20)=3.28, P=0.2) nor SLC17A7+ cells (Kruskal-Wallis ANOVA, H(2,21)=2.58, P=0.29). (**G**) Densities of PV+ cells assessed by immunohistology. No change between groups was observed (group effect: P=0.132; layer effect: P<0.0001; group x layer: P=0.083).

PNNs have been most extensively described around PV+ cells, but are also found around other neuronal types^22^. We first wanted to specify the identity of cells covered by PNNs in the human vmPFC. We measured the ratios of PNNs surrounding either PV+ cells, glutamatergic neurons, and other interneurons in a subset of control, psychiatrically healthy subjects (**Fig.1E-F**). Since PV antigenicity is particularly susceptible to freezing, and lost altogether in samples snap frozen prior to fixation, we developed an approach to combine fluorescence in situ hybridization (FISH) and immunofluorescence to visualize PVALB expressing cells and WFL+ PNNs in frozen samples. Similarly, glutamatergic neurons and other subtypes of interneurons were visualized with FISH using probes against SLC17A7 (vesicular glutamate transporter 1) and GAD1 (glutamate decarboxylase 1), respectively (**Fig.1E**). The majority of cells covered by WFL staining were PVALB+ (∼74%), followed by GAD1+/PVALB-cells (∼23%), indicating that a small fraction of PNNs is likely surrounding other subtypes of interneurons (**Fig.1F**). As previously reported^23,24^, some PNNs stained with WFL were also found to surround glutamatergic neurons (SLC17A7+), although only a very small fraction of them (∼3%, **Fig.1F**).

To clarify the cellular specificity of the observed increase in PNNs recruitment, we next quantified the ratios of PVALB+, SLC17A7+, and GAD1+/PVALB-cells surrounded by WFL+ PNNs. ∼65% of PVALB+ cells were surrounded by PNNs (**Fig.1G**), in line with previous observations^25^, while only a small proportion of GAD1+/PVALB- and SLC17A7+ cells were covered by PNNs (**Fig.1G**). Interestingly, ratios of cells covered by WFL-labeled PNNs showed a positive correlation with age (**Supplementary Table 1**), suggesting an increased recruitment of PNNs with age regardless of cell type. Importantly, samples from DS-CA individuals displayed a robust increase in the percentage of PVALB+ cells surrounded by PNNs compared to DS and CTRL samples (**Fig.1G)**, while no change in the proportion of other cell types covered by PNNs was found between group. Controlling for age as a covariate did not change the outcome of these group comparisons (%PVALB+/PNN+ cells: F(2,15)=4.08, P=0.047; %GAD1+/PVALB-/PNN+ cells: F(2,15)=0.237, P=0.793; %SLC17A7+/PNN+ cells: F(2,15)=1.430, P=0.281).

Finally, we addressed whether the increased densities of PNNs and higher ratios of PV+ cells surrounded by PNNs observed in DS-CA subjects could be linked to changes in the number of PV+ cells in the vmPFC and found no evidence of altered PV+ cells densities between groups (Fig.1H). Of note, PV+ cell densities were inversely correlated with age (**Supplementary Fig.1**), but controlling for this factor did not change the outcome of group comparisons (ANCOVA with group as fixed factor and age as covariate: group effect, P= 0.251; age effect, P=0.007).

Altogether these results suggest that a history of CA in depressed suicides is associated with increased recruitment and maturation/morphological complexity of PNNs around PV+ neurons, rather than changes in cell populations.

We then sought to indirectly explore the molecular underpinnings of this phenomenon and reasoned that increased recruitment of PNNs associated with CA should translate or be induced by changes in the molecular programs controlling PNN assembly. Our understanding of these transcriptional programs is scarce, hindered by the fact that several known molecules participating in PNN recruitment are released non-locally and by different cell types, implying a complex cellular crosstalk orchestrating PNN assembly. To gain insight into how, in humans, different cell types contribute to the synthesis of canonical components of PNNs, we explored a single-nucleus sequencing dataset previously generated by our group in the human dorsolateral PFC ^20^ (Brodmann area 9), and screened their expression across 8 major cell types. The main canonical components of PNNs, namely aggrecan (ACAN), neurocan (NCAN), versican (VCAN), phosphacan (PTPRZ1), brevican (BCAN), and tenascin-R (TNR), were highly enriched in oligodendrocyte progenitor cells (OPCs), in particular VCAN, PTPRZ1, BCAN and TNR (fold change of 140, 37, 7.9, and 22.9 respectively between gene expression in OPCs versus PV+ cells) (**Fig.2A and Supplementary Fig.1)**. Because this dataset originates from the dlPFC, and region-specific patterns of ECM-related gene expression could exist, this was further validated using FISH (**Fig.2B and 2C**) in the vmPFC for versican (VCAN) and phosphacan (PTPRZ1), as they showed the strongest expression in OPCs and are two major signature genes in late OPCs ^26^. We found that in the vmPFC, cells expressing these genes are almost all PDGFRA+ OPCs (97.9% of VCAN-expressing cells were co-expressing PDGFRA, and 92% of PTPRZ1-expressing cells were co-expressing PDGFRA) (**Fig.2D**). Interestingly, despite that VCAN gene expression was restricted to OPCs, immunolabeling of the versican protein showed a characteristic pattern of PNNs and an overlap with WFL-labeled PNNs (**Fig.2E**), suggesting overall that OPCs could be potent regulators of PNN formation.

**Figure 2.**
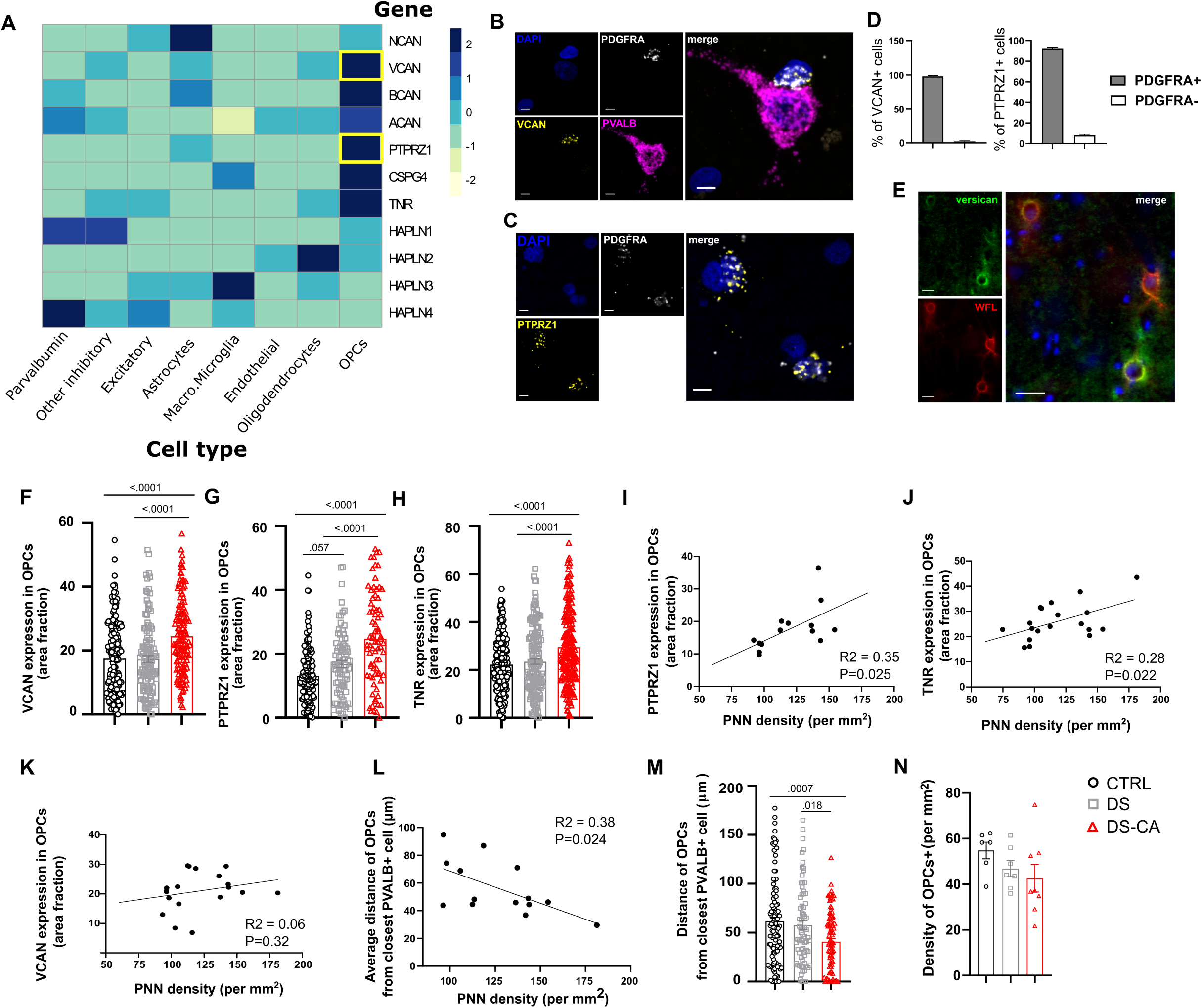
**(A)** Expression of canonical components of PNNs according to cell type, derived from single-nucleus RNA sequencing of 34 human dlPFC (BA9) samples ^20^. OPCs consistently express higher levels of most of these components compared to other cell types. (**B**) Representative images of FISH validation of VCAN (Versican, yellow) expression in OPCs (PDGFRA+ cells, white). Note the VCAN expressing OPC juxtaposed to a PVALB+ (magenta) cell. Nuclei were counterstained with DAPI (blue). Scale bar = 5 μm. (**C**) Representative images of FISH validation of PTPRZ1 (Phosphacan, yellow) expression in OPCs (PDGFRA+ cells, white). Nuclei were counterstained with DAPI (blue). Scale bar = 5 μm. (**D**) Both VCAN (left) and PTPRZ1 (right) expression is highly enriched in OPCs, with 97.8 percent of VCAN+ cells (N=225) co-expressing PDGFRA, and 91.8 percent of PTPRZ1+ cells (N= 281) co-expressing PDGFRA. (**E**) Representative image of versican (green) immunolabeling. Despite enrichment of VCAN gene in OPCs, the versican protein shows a characteristic PNN staining pattern and co-localized with WFL (red). Nuclei were counterstained with DAPI (blue). Scale bar = 25 μm. (**F**) The average expression of VCAN in OPCs was significantly higher in DS-CA subjects (N=139 cells, 7 subjects) compared to CTRLs (N= 160 cells, 8 subjects) and DS (N= 119 cells, 6 subjects) (One-way ANOVA F (2, 415) = 17.25, P<0.0001, followed by Tukey’s multiple comparison test). (**G**) The average expression of PTPRZ1 in OPCs was significantly higher in DS-CA subjects (N=63 cells, 4 subjects) compared to CTRLs (N= 117 cells, 6 subjects) and DS (N= 81 cells, 5 subjects) (One-way ANOVA F (2, 258) = 31.65, P<0.0001, followed by Tukey’s multiple comparison test). (**H**) The average expression of TNR in OPCs was significantly higher in DS-CA subjects (N=200 cells, 8 subjects) compared to CTRLs (N= 207 cells, 7 subjects) and DS (N= 160 cells, 5 subjects) (One-way ANOVA, F (2, 564) = 18.69, P<0.0001, followed by Tukey’s multiple comparison test). Both PTPRZ1 (**I**) and TNR (**J**), but not VCAN (**K**) average expression in OPCs modestly correlated with PNNs densities (R^2^ = 0.35, P=0.025 and R^2^ = 0.28, P=0.022 respectively). (**L**) A negative correlation was found between average distance of OPCs from closest PVALB+ cell and PNNs density (R^2^ = 0.36, P=0.024), suggesting that OPCs proximity with PVALB+ cells could be associated with changes in PNN density. (**M**) Proximity of OPCs with PVALB+ cells was increased in DS-CA subjects (N=90 OPCs, 5 subjects) compared to CTRLs (N= 106 OPCs, 6 subjects) and DS (N= 73 OPCs, 4 subjects) (One-way ANOVA: F (2, 266) = 7.963, P=0.0004, followed by Tukey’s multiple comparison test). (**N**) Average densities of PDGFRA+ OPCs were not changed between DS-CA (N=8), DS (N=7) and CTRL (N=6) groups (Kruskal-Wallis ANOVA, H(2,21)=4.67, P=0.095).

In support of a possible involvement of OPCs in mediating CA-related changes in PNNs, the expression of VCAN, PTPRZ1 and TNR was upregulated in OPCs of CA victims (**Fig.2F-H**), and both the expression of PTPRZ1 and TNR in OPCs correlated with WFL-labeled PNNs densities regardless of group (**Fig.2I and 2J**).

OPCs were on occasion directly juxtaposed to PVALB+ cells (**Fig.2B**), as previously reported in rodents ^27^. Interestingly, OPC proximity to PVALB+ cells modestly correlated with PNN density (**Fig.2L**, R2 = 0.36, P=0.024), and was increased in individuals with a history of CA (**Fig.2M**), further suggesting an interplay between these two cell types. To clarify if these changes could be linked or associated with changes in cell numbers, we compared the density of OPCs between groups. Interestingly OPC densities showed a marked decrease with age (**Supplementary Table 1**), but no difference between groups were found (**Fig.2N**), even after controlling for this factor (ANCOVA, with group as fixed factor and age as covariate: group effect, P= 0.13; age affect, P=0.001).

## Discussion

Overall, our results suggest that a history of CA may associate with increased recruitment and maturation of PNNs, as well as an upregulation of their canonical components by OPCs, a cell type that likely plays a key role in the cellular crosstalk that orchestrates PNN formation.

Although, to our knowledge, this is the first evidence in humans that early-life adversity (ELA) affects the recruitment of PNNs, recent studies in animals have approached this question. Guadagno et al.^28^ found that in the limited bedding paradigm, adolescent pups have increased densities of PNNs in the amygdala. Murthy and al.^9^ showed that in the ventral dentate gyrus, maternal separation combined with early-weaning, another model of early-life stress mimicking some aspects of adversity, led to an increase in PNN intensity around parvalbumin-positive interneurons with no change in PNN density. Importantly, these effects were present in adults, suggesting a long-lasting impact of ELA on PV+ cell function and PNN remodeling. Gildawie et al.^29^ also recently reported that in the prelimbic cortex, maternal separation in rats increased the intensity of PNNs surrounding PV+ neurons, an effect only observed in adulthood. This suggests that changes in PNN integrity and maturation following ELA could possibly be protracted and develop over time. While we can’t address this question with our post-mortem design, our results show at the very least that changes in PNN integrity in victims of CA are observable at adult age. Few studies have so far been conducted on this topic, however, with one of them showing a decrease in the intensity of WFA fluorescent labeling in both the prefrontal cortex and hippocampal CA1 following an early-life sub-chronic variable stress paradigm^30^. Clearly, more work is needed to better characterize the effects of ELA on PNNs integrity, particularly in light of the multiple paradigms used in animal studies.

Despite the fact that much of the literature has focused on the influence of PNNs on PV+ cell physiology, it is important to note that PNNs are not exclusively present around PV+ neurons^22,23,31^. Although our data indicate that in the vmPFC PV-surrounding PNNs are the majority, a portion of them were identified around other GABAergic neurons, as well as around a small fraction of excitatory neurons. Our data suggest that CA associates with a selective increase in the recruitment of PNNs around PV+ neurons, considering that the higher percentage of cells surrounded by WFL-labeled PNNs was only found for PV+ cells, but not for other cell types. However, we cannot exclude that given the smaller pool of non-PV cells surrounded by WFL-labeled PNNs, we were unable to detect such effects in our design. It is also likely that the sole use of WFL immunostaining to detect PNNs may be a limitation to fully understand their distribution. It is becoming increasingly clear that PNNs vary in molecular composition, and perhaps function, and that WFL-labeled PNNs may be biased towards specific neuronal types^32,33^. The use of additional markers should help decipher how ELA affects the remodeling of the ECM more broadly. It is also noteworthy that this analysis, by focusing on cortical layers IV-V, did not allow to clarify the possible layer specificity of PNN cellular distribution. Given the molecular, cellular, and connectivity heterogeneity in different cortical layers, it is possible that PNNs differentially interact with these different cell types and that ELA may affect these interactions in a layer specific manner.

One particularly noteworthy aspect of our results is the specific association between changes in PNNs and a history of CA. When comparing DS and controls for the expression of all PNN-related genes reported in our study (Figure 2A), based on the snRNAseq data generated by Nagy et al.^20^ (Supplementary Tables 30-31), none showed differential expression between groups in OPCs. The fact that our FISH experiments did not show significantly enhanced expression of these markers in DS samples is therefore consistent both with results obtained by Nagy et al. and with our own findings indicating that PNNs are more abundant and mature specifically in samples from DS with a history of CA. Overall, this suggests that while transcriptomic changes in OPCs may be a strong feature of depressed suicides^20^, PNN-related changes are more specific to a history of CA. Because such changes are absent in depressed suicides without a history of CA, our findings suggest possible vulnerability windows during which PV+ cell function and PNN maturation are more susceptible to experience-dependent remodeling and adversity. If these changes may mediate some of the negative mental health outcomes or cognitive and emotional traits associated with CA in adulthood, they likely don’t represent a hallmark of depression, in accordance with a recent post-mortem study finding no change in the density of PNNs in the PFC of depressed patients^34^.

The search for possible mechanisms involved in the effects of ELA on PNNs development is an unexplored field. PNN recruitment is likely orchestrated by a complex interplay between activity-dependent autonomous pathways in parvalbumin neurons, with signals originating from different cell types involved in their assembly. Because parvalbumin neurons have been shown to be particularly sensitive to stress and glucocorticoids, in particular early-life stress^35–40^, we can speculate that elevated glucocorticoids in CA victims^41^ can impact PV+ neuron function early-on. This could translate into increased GABA release following GR activation^39^, and indirect increase in the recruitment of PNNs, which has been directly linked to PV+ neuron activity^42,43^ and GABA levels^44^. A myriad of factors could however indirectly affect PV+ cells during this period of protracted maturation associated with childhood and adolescence, such as increased pro-inflammatory cytokine expression associated with ELA^36,45^ or changes in neurotrophic factor expression^46–48^.

An interesting molecular candidate is the transcription factor OTX2, released non-locally by cells in the choroid plexus, and acting as a major initiator of PNN development^49^. Recent evidence suggests a role of OTX2 in mediating vulnerability to early-life stress^50^, and Murthy et al. reported elevated expression of OTX2 in the choroid plexus following maternal separation and around PV+/PNNs+ cells in the ventral dentate gyrus^9^. While the precise mechanisms involved in the effects of ELA on the release of OTX2 are not known, it is noteworthy that DNA methylation of the OTX2 gene in children has been shown to correlate with increased risk for depression as well as increased functional connectivity between the vmPFC and bilateral regions of the medial frontal cortex^51^. This highlights that beyond discrete changes in PV+ cell function, ELA could affect the release of distal cues by non-neuronal cells and contribute to extracellular matrix remodeling, thus affecting brain function and vulnerability to psychopathology.

As mentioned previously, although PNN development has been strongly linked to neuronal activity^3,42,52^, PNN integrity and assembly are likely orchestrated by the complex integration by PV+ neurons of cues originating from multiple cell types^53,54^. Accordingly, we found that the expression of genes encoding for the major canonical components of PNNs were strongly enriched in oligodendrocyte-lineage cells, in particular in OPCs, while PV+ neurons barely expressed any of those components. While our single-cell expression data originates from the dorsolateral PFC, this was validated in the vmPFC, thus decreasing the possibility that this pattern of enrichment is region-specific. This is also in accordance with previous literature in rats, albeit in the cerebellum, similarly showing that VCAN, PTPRZ1 and TNR are almost exclusively expressed in oligodendrocyte-lineage cells^55^. This is also consistent with more recent single-cell RNAseq studies showing an enrichment of extracellular components, including VCAN and PTPRZ1, in OPCs^56–58^.

The interplay between oligodendrocyte-lineage cells and PV+ neurons, in particular during developmental windows of plasticity, are being increasingly documented^59^. Interestingly, OPCs have been shown to be ontogenetically related to PV+ neurons^60^. OPCs also receive functional synaptic inputs from GABAergic interneurons, a connectivity that reaches its peak during early postnatal development along with the maturation of PNNs^61–63^. It is therefore particularly tempting to speculate that OPC-PV neuron communication during critical windows of development may play a fundamental role in modeling cortical plasticity and the maturation of PNNs. This has never been addressed, and we therefore approached this question by investigating how CA associates with changes in the expression of PNN canonical components specifically in OPCs, and how this correlates with the changes in PNN integrity observed in victims of CA. First, we found that OPC proximity to PV+ neurons positively correlated with PNN densities, and that in victims of CA, OPCs tended to be more proximal to PV+ neurons. Since a proportion of OPCs are in direct physical contact with neuronal populations, and preferentially around GABAergic neurons^27,64^, we could speculate that CA associates with increased recruitment of OPCs around PV+ cells, thereby promoting the formation of PNNs around these cells. Caution is however needed in interpreting these correlational results that functionally relate the distance between OPCs and PV+ cells to the recruitment of PNNs. The high percentage of PV+ cells enwrapped by PNNs in our data (∼65%) and low percentage of PV+ cells paired with an OPC (∼3% based on Boulanger and Tessier, 2017, albeit in rodents) rather indicates that is unlikely that direct pairing of OPCs is necessary for the formation of PNNs. This question should therefore be further addressed with appropriate functional tools in cellular or animal models. Of important note, we did not address whether OPCs proximity with other cell types was changed, and how it could relate to changes in various cell populations. Additionally, while we did not observe any change in overall OPCs densities between groups, a more refined layer-specific analysis of OPCs densities would be informative to rule out the influence of cell numbers on the physical proximity of these two cell types.

Secondly, we also found that the expression of canonical components of PNNs in OPCs was positively correlated with PNN densities and upregulated in victims of CA. This strongly suggests that CA has a durable impact on OPC molecular programs that may contribute to PNN development. While we focused on VCAN, PTPRZ1 and TNR based on their high enrichment in OPCs, it would be important to describe more broadly how CA affects the transcriptional signature of PNNs. In particular, some of these components, while present in PNNs, are also found in other ECM compartments, such as the perinodal ECM and around excitatory cells, contributing to synaptic function^3^. While we did not address this question, the impact of CA on OPC function may therefore more broadly affect ECM physiology and remodeling. It is also becoming increasingly clear that multiple populations with distinct functional or transcriptomic features are encompassed in OPCs^65,66^. While we can speculate that these subtypes may have distinct role in the remodeling of the ECM, we did not address this important question.

Our results, nonetheless, further strengthen previous reports by our group^67,68^ that CA has profound effects on oligodendrocyte-lineage cells, that may extend well beyond affecting myelination by contributing to the reprogramming of various aspects of brain plasticity.

How precisely OPC-PV communication contributes to the development of PNNs and impacts PV+ neurons functional integration and circuit dynamics certainly deserves further investigation using functional approaches. Inherent to our post-mortem design, a major limitation of this study lies in our inability to infer on the precise timing of these changes, and whether dynamic remodeling of the extracellular matrix occurs in a protracted way. Given the correlational nature of our results, we cannot infer on the possible influence of recent and emotionally salient events on PNN remodeling, or possible state-dependent factors at the time of death. Several rodent studies have indeed reported that behavioural manipulations in adulthood, in particular learning and memory paradigms in which plasticity events are recruited, can affect PNN dynamics^69–72^.

While it is also tempting to infer that CA selectively affects PNN remodeling around PV+ neurons, caution is needed in this interpretation. As previously mentioned, WFL immunostaining may only label a specific fraction of PNNs^32,33^, and the continuum between PNN components and other ECM compartments imply more refined approaches are necessary to fully understand how CA impacts ECM remodeling and the role of OPCs in this form of plasticity.

Another limitation is that our study only included very few female samples, given the much higher prevalence of suicide in males. Sex is known to moderate both the biological and the psychopathological effects of CA^73^, and increasing evidence points towards a sexual dimorphism in the effects of stress, in particular early-life stress, on PV+ neuron function and perhaps PNN development^8,40,74^. Future studies should therefore explore this aspect to further understand how ELA modifies trajectories of brain development.

Other limitations are inherent to post-mortem studies of psychiatric cohorts, such as presence of medication in some subjects. Although this was the case for both depressed suicides groups, and that the observed changes seem specific to a history of CA, we cannot exclude interactive effects of medical treatments and life trajectories on our variables. Similarly, relatively long PMIs in our cohorts, although not statistically linked to changes in our variables, could potentially confound our results.

To conclude, our findings suggest that CA may lead to persistent patterns of maladaptive behaviors by reducing the neuroplasticity of cortical circuits through the enhancement of developmental OPC-mediated PNN formation. Future preclinical models should help determine whether changes in OPCs are causal in the increased recruitment of PNNs following CA or an indirect response following altered PNN dynamics. Likewise, the consequences of these molecular changes should be examined at the network level to determine their functional impact on intra- and inter-regional communication.

## Acknowledgments and funding

This work was funded by a CIHR Project grant (PJT-173287) to NM. AT was supported by fellowships from the FRQS and Toronto Dominion, and an American Foundation for Suicide Prevention (AFSP) Young Investigator Innovation Grant (YIG-0-146-17). The Molecular and Cellular Microscopy Platform and the Douglas-Bell Canada Brain Bank (DBCBB) are partly funded by a Healthy Brains for Healthy Lives (CFREF) Platform Grant to GT and NM. The DBCBB is also funded by the Réseau Québécois sur le suicide, le troubles de l’humeur et les troubles associés (FRQS).

## Author Contributions

AT and NM conceived the study. GT participated in the acquisition and clinical characterization of the brain samples. AT, CB, FD, MAD, CC, RM, AM contributed to immunohistological experiments. AT, CN, MM and KP generated and analyzed the snRNA-seq dataset. AT, CB, FC, ST performed the in-situ hybridization experiments. AT, CB and NM prepared the manuscript and all authors contributed to and approved its final version.

