## Supplementary figures and images for "Child abuse associates with increased recruitment of perineuronal nets in the ventromedial prefrontal cortex: a possible implication of oligodendrocyte progenitor cells"

### Supplemental Figure 1

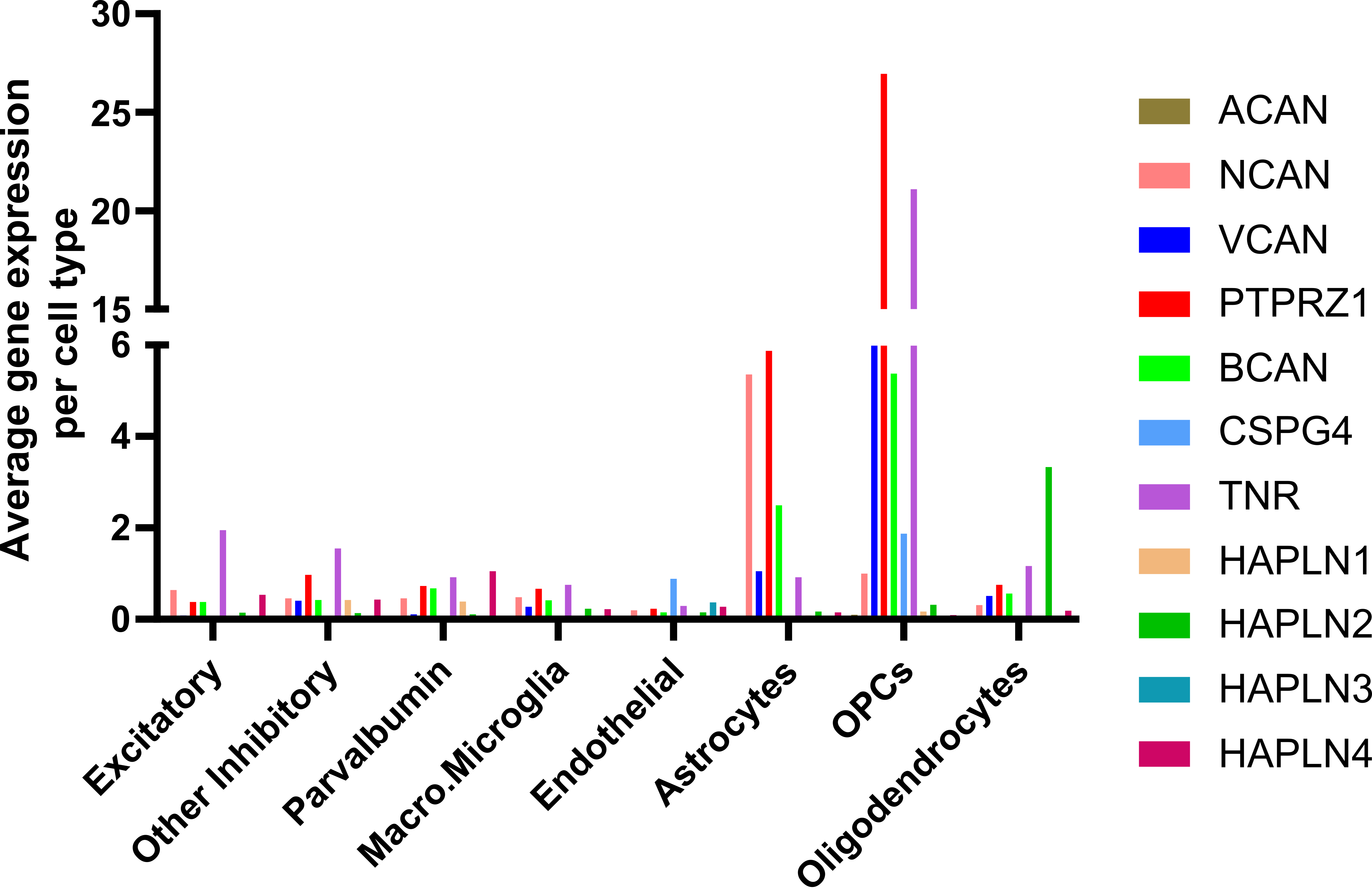
