## Supplemental Table 1 for "Child abuse associates with increased recruitment of perineuronal nets in the ventromedial prefrontal cortex: a possible implication of oligodendrocyte progenitor cells"

| Spearman’s rho |  | PNNs densities | WFL intensity per PNN | coverage per PNN | PV+ cells densities | %PVALB+ cells covered by PNNs | %SLC17A7+ cells covered by PNNs | %GAD1+/PVALB- cells covered by PNNs | PTPRZ1 expression in OPCs | VCAN expression in OPCs | TNR expression in OPCs | PDGFRA+ (OPCs) cells densities |
| --- | --- | --- | --- | --- | --- | --- | --- | --- | --- | --- | --- | --- |
| Age | Coef | .042 | .156 | .015 | -.411^**^ | .433^*^ | .472^*^ | .586^**^ | .258 | -.078 | -.131 | -.669^**^ |
|  | Sig. | .810 | .594 | .958 | .007 | .050 | .035 | .007 | .354 | .737 | .582 | .001 |
| PMI | Coef | -.008 | -.349 | .165 | .241 | .231 | .006 | -.147 | -.306 | .263 | -.094 | -.351 |
|  | Sig. | .964 | .221 | .573 | .124 | .313 | .980 | .535 | .268 | .250 | .693 | .119 |
| pH | Coef | .115 | .033 | -.145 | .038 | .220 | .113 | -.021 | .018 | -.153 | -.185 | -.387 |
|  | Sig. | .511 | .911 | .620 | .811 | .337 | .635 | .930 | .949 | .507 | .434 | .083 |
| Substance dependence | Coef | -.101 | -.194 | -.324 | .100 | .280 | .272 | .070 | .091 | .280 | .369 | .045 |
|  | Sig. | .564 | .506 | .259 | .527 | .218 | .245 | .769 | .748 | .218 | .110 | .847 |
| Medication | Coef | .058 | -.075 | .447 | -.114 | .140 | .130 | .338 | .194 | -.393 | .392 | -.196 |
|  | Sig. | .774 | .828 | .168 | .535 | .620 | .658 | .259 | .568 | .148 | .165 | .483 |

**Supplementary Table 1.** Correlation coefficients and P values for Spearman’s rho non-parametric measure of association between co-variates and dependent variables.
